# Selective increase of functional network connectivity in Arc-positive neuronal engrams after long-term potentiation

**DOI:** 10.1101/2020.12.07.415109

**Authors:** Yuheng Jiang, Antonius M.J. VanDongen

## Abstract

New tools in optogenetics and molecular biology have culminated in recent studies which mark immediate-early gene (IEG)-expressing neurons as memory traces or engrams. Although the activity-dependent expression of IEGs has been successfully utilised to label memory traces, their roles in engram specification is incompletely understood. Outstanding questions remain as to whether expression of IEGs can interplay with network properties such as functional connectivity and also if neurons expressing different IEGs are functionally distinct. We investigated the expression of Arc and c-Fos, two commonly utilised IEGs in memory engram specification, in cultured hippocampal neurons. After pharmacological induction of long-term potentiation (LTP) in the network, we noted an emergent network property of refinement in functional connectivity between neurons, characterized by a global down-regulation of network connectivity, together with strengthening of specific connections. Subsequently, we show that Arc expression correlates with the effects of network refinement, with Arc-positive neurons being selectively strengthened. Arc positive neurons were also found to be located in closer physical proximity to each other in the network. While the expression pattern of IEGs c-Fos and Arc strongly overlaps, Arc was more selectively expressed than c-Fos. These IEGs also act together in coding information about connection strength pruning. These results demonstrate important links between IEG expression and network connectivity, which serve to bridge the gap between cellular correlates and network effects in learning and memory.

## INTRODUCTION

Activity-driven neuronal plasticity is the basis for learning and memory. New tools in optogenetics and molecular biology have culminated in numerous studies in recent years which mark immediate-early gene (IEG)-expressing neurons as memory engrams (Liu 2012, Denny 2014, Tonegawa 2015). Transcriptional programs activated upon neuronal stimulation are complex, with the involvement of a myriad of IEGs, their downstream effectors and the interplay between gene transcription and ongoing neuronal activity (Yap 2018). Although the activity-dependent expression of IEGs have been successfully utilised to label memory traces, their roles in engram specification is incompletely understood. For instance, although the expression profiles of IEGs within neuronal populations overlap significantly (Mahringer 2019), they are diverse and distinct (Gonzales 2019), with different IEGs performing distinct functions. Furthermore, it is known that within single neurons, there is a combinatorial expression of IEGs (Sheng 1995). This suggests that differential expression of IEGs can encode the specificity of the stimulus, which has been demonstrated in part using single-cell RNA-sequencing technology (Jaeger 2018). Arc/Arg3.1 (activity-regulated cytoskeleton-associated protein/activity-regulated gene 3.1) and c-Fos are two IEGs that are commonly used in memory engram labelling experiments (Kawashima 2014). Arc is an important player in synaptic plasticity and long-term memory consolidation (Plath 2006, Shepherd 2011, Nikolaienko 2018). c-Fos is another IEG whose increase in expression is commonly used as an indicator for neuronal activation. It was one of the first transcription factors whose induction was shown to be activity-dependent (Sheng 1990). It has been recently shown that IEGs Arc and c-Fos are differentially expressed within neuronal populations and their combinatorial expression is functionally linked to whether a memory trace is newly formed or reactivated (Jaeger 2018). This is due to the difference in the expression dynamics of the two IEGs, with c-Fos being transiently expressed after activation but Arc maintaining a high level of transcription hours after stimulus onset. Further questions remain, however, as to whether expression of IEGs can interplay with network properties such as functional connectivity and also if neurons expressing different IEGs are functionally distinct.

The fundamental cellular correlate of memory is synaptic plasticity, while engram formation is dependent on the connectivity of the neuronal network (Tonegawa 2015). Conventionally, studies of neuronal activity and IEG expression are mostly based on the single cell level whereas network activity is recorded on the neuronal population level, thus there is a lack in linking IEG expression and changes in plasticity on the cellular level with changes in network responses. Similarly, studies on long-term potentiation (LTP) are extensive on the molecular, cellular and pathway level, but few studies have focused on the effects of LTP in altering network responses. Among the various induction methods of LTP, direct chemical stimulation of specific biochemical processes to induce LTP can result in plasticity changes in large numbers of synapses from multiple neurons. NMDA receptor dependent LTP requires the coincidence of presynaptic glutamate release and postsynaptic depolarisation. Chemically, this can be accomplished by a variety of pharmacological agents. Among these, bicuculline, a GABAA receptor antagonist has been shown to be able to induce bursts of action potential and elicit NMDA receptor-dependent calcium transients. When this is applied together with the presynaptic K^+^ channel blocker 4-amino pyridine (4-AP), which enhances glutamate release, calcium transients in the neuronal nucleus increase from baseline levels (Hardingham 2001). This sustained elevation in nuclear calcium can increase the affinity of calmodulin binding to calcium and in turn result in maintained activation of Ca^2+^/calmodulin-dependent enzymes that potentially facilitate transcription. The synergistic effects of these agents have been shown to be able to increase the endogenous expression of cFos (Hardingham 2001). Forskolin, an adenylyl cyclase activator, can effectively increase the level of cAMP levels and directly activate transcription and translation that is important for the expression and maintenance of protein synthesis-dependent late phase of LTP (Otmakhov 2004). It has also been shown that the translational repression of the IEG Arc is alleviated by application of forskolin, through activation of the cAMP-dependent protein kinase (PKA) pathway (Bloomer 2008). This combination of pharmacological agents, namely 4-AP, bicuculline and forskolin, has been used effectively to induce LTP in cultured hippocampal neurons in the study of Arc dynamics (Wee 2014, Oey 2015, Leung 2019).

To study interplay between IEG expression and network activity, we made use of calcium imaging, and changes in calcium levels as a proxy for activity changes. Changes in functional network connectivity can be studied systematically (Poli 2016), and the results for individual neurons can be mapped onto and correlated with the expression of Arc and c-Fos in these neurons. In so doing, we aim to address important links between IEG expression and network connectivity, which serve to bridge the gap between cellular correlates and network effects in learning and memory.

## MATERIALS AND METHODS

### Neuronal Cell Culture

Hippocampi and/or cortices from embryonic day 18 Sprague-Dawley rats of either sex was dissected in ice-cold Hanks Balanced Salt Solution (HBSS) and digested using a papain dissociation system (Worthington Biochemical Corporation). The appropriate density of neurons was plated on glass-bottom culture dishes (MatTek, Ashland, MA) that had been coated with poly-D-lysine at 0.1mg/ml (Invitrogen) for 2 hours at 37 degrees. Neurons were cultured in Neurobasal medium supplemented with 2% B-27, 0.5 mM L-glutamine, 10% penicillin/streptomycin, and half of the medium is replaced bi-weekly from day-in-vitro (DIV) 5 onwards. All animal procedures were performed in accordance with the Institutional Animal Care and Use Committee (IACUC) regulations.

### Neuronal Stimulation and Silencing

We used a combination of 4-Aminopyridine (4AP), Bicuculline (Bic), and Forskolin at final concentrations of 100μM, 50μM, and 50μM respectively (4BF) to stimulate synaptic NMDA receptors and network activity and chemically induce LTP. The drugs were added to the culture medium for two to four hours. Activity silencing was done by adding tetrodotoxin (TTX) at a concentration of 1uM to the culture medium before treating the culture with the stimulation drugs.

### Conventional Immunofluorescence

Cells were fixed with a solution containing 4% paraformaldehyde (PFA), 4% sucrose, and PBS for 15 minutes at room temperature (RT), blocked with a solution containing 10% goat serum, 2% bovine serum albumin (BSA), and PBS for 1 hour at RT. Rabbit-anti-NeuN (Thermo Fisher Scientific Cat# 711054, RRID:AB_2610583), mouse-anti-Arc (C7) (Santa Cruz Biotechnology Cat# sc-17839, RRID: AB_626696) and rabbit-anti-c-Fos (Santa Cruz Biotechnology Cat# sc-52, RRID:AB_2106783) primary antibody were used at 1:300. Primary antibodies were incubated for 1 hour at RT in a dilution buffer containing 1:1 block solution and PBS-Triton X solution at 1:300. Dishes were washed 3 times with PBS-Triton X and incubated with Alexa-Fluor 488 conjugated secondary antibody 1:1000 (Molecular Probes-Invitrogen) in dilution buffer for 1 hour at RT. Washing was repeated as per the above. The cells were subsequently incubated with 5μM DAPI for 5 minutes before being imaged.

### Transduction

For calcium imaging, the GCaMP6m construct (pAAV.Syn.GCaMP6m.WPRE.SV40) was a gift from Douglas Kim & GENIE Project (Addgene viral prep # 100841-AAV9); http://n2t.net/addgene:100841; RRID:Addgene_100841), and transferred from University of Pennsylvania, Vector Core. It was transduced into the cells at a MOI of 1 × 10^5^ on DIV8. Half of the medium was changed on DIV9 to prevent virus toxicity.

### Widefield Imaging

Fluorescence images were obtained using a motorized inverted wide-field epifluorescence microscope (Nikon Eclipse Ti-E), using Nikon 10X Plan Apo objective (N.A. = 0.4). Motorized excitation and emission filter wheels (Ludl electronics, Hawthorne, NY) fitted with a DAPI/CFP/YFP/DsRed quad filter set (#86010, Chroma, Rockingham, VT) were used together with filter cubes for DAPI, YFP and TxRed (Chroma) to select specific fluorescence signals.

### Calcium imaging

Calcium imaging using widefield microscopy was done on neuronal cultures transduced with GCaMP6. Movies of 1-minute duration were taken at a frame rate of 5Hz at baseline, after 4BF treatment and post 4BF removal. Upon completion of calcium imaging, the culture was fixed and stained as described. Fig. 1A shows the calcium imaging and analysis pipeline.

**Figure 1.**
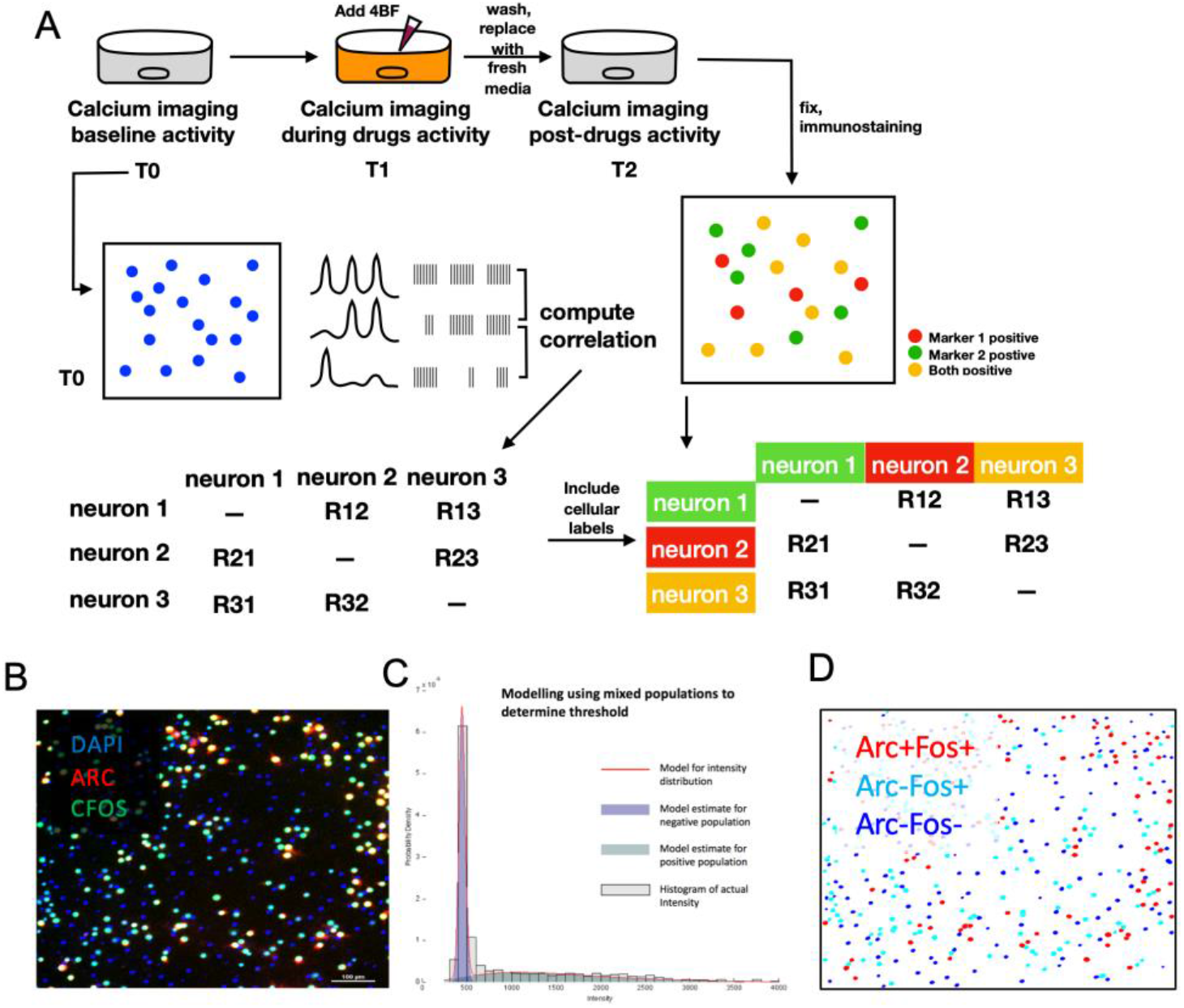
Calcium imaging pipeline and data analysis. **A)** Schematic of calcium imaging: cells transduced with GCaMP6 are imaged under widefield imaging at baseline and after 4BF treatment. Calcium movies are processed to extract traces of individual neurons, which are converted into spike times using methods described. The correlation between neuronal spike times is calculated using the spike time tiling coefficient (STTC). Cells were fixed and stained with the desired markers (NeuN with Arc or Arc with c-Fos). These were then used as identity markers for subsequent connectivity analysis. **B)** Representative immunofluorescence staining image showing Arc and c-Fos with DAPI. Extraction of calcium waves of individual neurons are based on selection of regions of interest (ROIs) by segmentation of the DAPI signals. **C)** Neuronal ROIs are determined with NeuN staining, and the threshold is selected based on mixed population modelling. The figure here shows the separation of staining intensities into a positive population and a negative one. A similar method is used to determine the threshold for Arc-positive and c-Fos-positive neuronal populations. **D)** Representative post analysis reconstruction image indicating ROI determination by DAPI and allocation of ROIs according to Arc and c-Fos identity as determined by their expression levels. These ROIs were subsequently used to extract calcium levels from within the ROI for subsequent spike time conversion and analysis.

### Data analysis

All custom written code used for analysis are available on GitHub at https://github.com/yuhengj/network_arc_dynamics.

### Calcium transient and spike detection

Images were acquired with the Nikon NIS Elements software and subsequently processed with MATLAB, with the BioFormats plugin. Calcium transients from the whole network were computed based on the total fluorescence change over time. Photobleaching was accounted for with fluorescence intensity from cells that were silent during the recording (lowest tenth percentile of the whole population of cells). Peaks in fluorescence were detected using MATLAB function *findpeaks.* To obtain calcium signals from only the neuronal population, regions of interest (ROIs) were first determined from the DAPI images, and the expression levels of NeuN within the ROIs were quantified (Fig. 1B). NeuN positive threshold was determined by mixed population modelling with expression levels from all cells (Fig. 1C), and the threshold for negative population was selected to be 3 standard deviations from the population mean. These NeuN positive ROIs are then used to extract fluorescence signals from the calcium movies recorded. All pixels within each ROI are averaged to give a single time course, and calcium transients (ΔF/F) were calculated by subtracting each value with the mean of signals from the lowest 10% of neurons at each time point. This removes background and corrects for photobleaching over time. Spike times were then inferred from ΔF/F using a maximum *a posteriori* (MAP) estimate of the spike train developed by Joshua Vogelstein (Vogelstein 2010) named fast-oopsi. This implementation was adapted to be used in MATLAB as part of the FluoroSNNAP package developed by Patel and colleagues (Patel 2015), which is available for download at https://www.seas.upenn.edu/~molneuro/software.html. Firing rates of neurons were calculated as the number of spikes divided by the total recording time.

### Functional connectivity

Neuronal correlations were calculated using the spike time tiling coefficient (STTC) that was previously described (Cutts 2014). This was chosen as this measure is known to be independent of firing rate. Specifically, the index of correlation is calculated according to the equation (1) below:

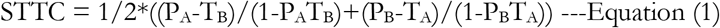

where T_A_ is the proportion of total recording time which lies within plus minus delta t of any spike from spike times of A. P_A_ is the proportion of spikes from A which lie within a small time window (±Δt) of any spike from B. Δt was set to 50 ms according to the previous study (Cutts 2014). T_B_ and P_B_ are calculated correspondingly, for any two given spike trains A and B. A maximum value of +1 can be obtained for complete correlation. This pair-wise synchronization index for each neuron can be used to populate a connectivity matrix M of size n × n, where n = number of neurons in the network. We used functions from the Brain Connectivity Toolbox (available for download at https://sites.google.com/site/bctnet) to edit the connectivity matrix.

### Change in network connectivity

To determine the change in connectivity of the whole network, we borrowed a measure developed for functional MRI that can represent all the connection strengths in the network, termed the Intrinsic Connectivity Distribution (ICD) (Scheinost 2012). This metric allows all of the connectivity information to be captured with only a few parameters and eliminates the need for setting arbitrary thresholds. ICD models the distribution of the connection strength first as a degree metric, from which a survival curve can be obtained and modelled as a stretched exponential with variance parameter (α) and shape parameter (β) according to equation (2),

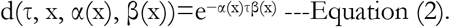

These parameters allow for a comparison of the connectivity distribution within a network, without the need for setting any arbitrary threshold for connectivity and enables comparison of a network across different timepoints.

### Statistical analysis

Statistical analysis was conducted using GraphPad Prism Software (La Jolla, CA). Data are presented as mean with SEM and statistical significance was set at *P<0.05, **P <0.01, ***P<0.001, ****P<0.0001.

### Regression model for Arc positive percentage

The *stepwiselm* function from MATLAB was used to find optimal terms from a list of firing rates (initial, final and change), connectivity (initial, final and change), refinement status (binary) to explain the percentage of Arc. The statistics of the regression model was subsequently obtained.

### Potentiated and depressed connections

Potentiated connections were defined as those that have initial connectivity indices between the values of 0 and 0.5 and final values between 0.5 and 1 while depressed connections are defined as those that are initially weak between 0 and 0.5 and have final values between 0.5 and 1. The corresponding value of the percentage of Arc-positive to Arc-positive connections is determined by counting the numbers of Arc-Arc connections over the total number of connections at each connectivity threshold. For potentiated connections, the threshold is based on the final connectivity index while for depressed connections the thresholds are determined based on the initial connectivity index.

### Distance measure

Each neuronal nuclear ROI from which the immunofluorescent signal and calcium signal is extracted from, has a central coordinate (x and y coordinates based on pixels). Each type of connection between neuronal pairs therefore corresponds to a physical Euclidean distance that can be calculated based on the central coordinate. Each image frame was 1007 × 1007 pixels, with each pixel corresponding to 0.8 μm, giving a total area of 0.64 mm^2^.

## RESULTS

To achieve LTP in cultured hippocampal neurons, we made use of a combination of 4-aminopyridine, a blocker of presynaptic K_v_1 family K^+^ channels, bicuculline, a γ-aminobutyric acid (GABA) receptor antagonist, together with forskolin, an adenylyl cyclase activator which enhances NMDA-dependent LTP (Otmakhov 2004). This cocktail, subsequently referred to as 4BF, increased synchronised network bursting in hippocampal cultures. Calcium imaging and analysis were conducted as described in the Materials and Methods section and outlined in Fig. 1A.

### 4BF increases neuronal firing rate but decreases overall connectivity

To assess the effects of 4BF on individual neurons, calcium traces for single neurons were extracted from the total recording. A representative raster plot of the neuronal firing at baseline, during 4BF treatment and after 4BF treatment can be seen in Fig. 2A. We found that there was a significant increase in the firing rate in neurons after 4BF treatment (Fig. 2A, B). To determine the functional connectivity between different neurons in the culture, the spike time tiling coefficient (STTC) was used to determine the index of correlation between all neuronal pairs (Cutts 2014). This revealed differences in initial culture states, with approximately 30% the cultures (n = 8 out of total 28) being well-connected prior to the treatment by 4BF and the remaining dishes being low in connectivity initially. Representative images of the connectivity between neurons are represented as heatmaps in Fig. 3A & B, with a highly connected initial state (Fig. 3B, first panel) or a less well-connected initial state (Fig. 3A, first panel). The average STTC for each network was found to be decreased as compared to baseline (Fig. 2C).

**Figure 2.**
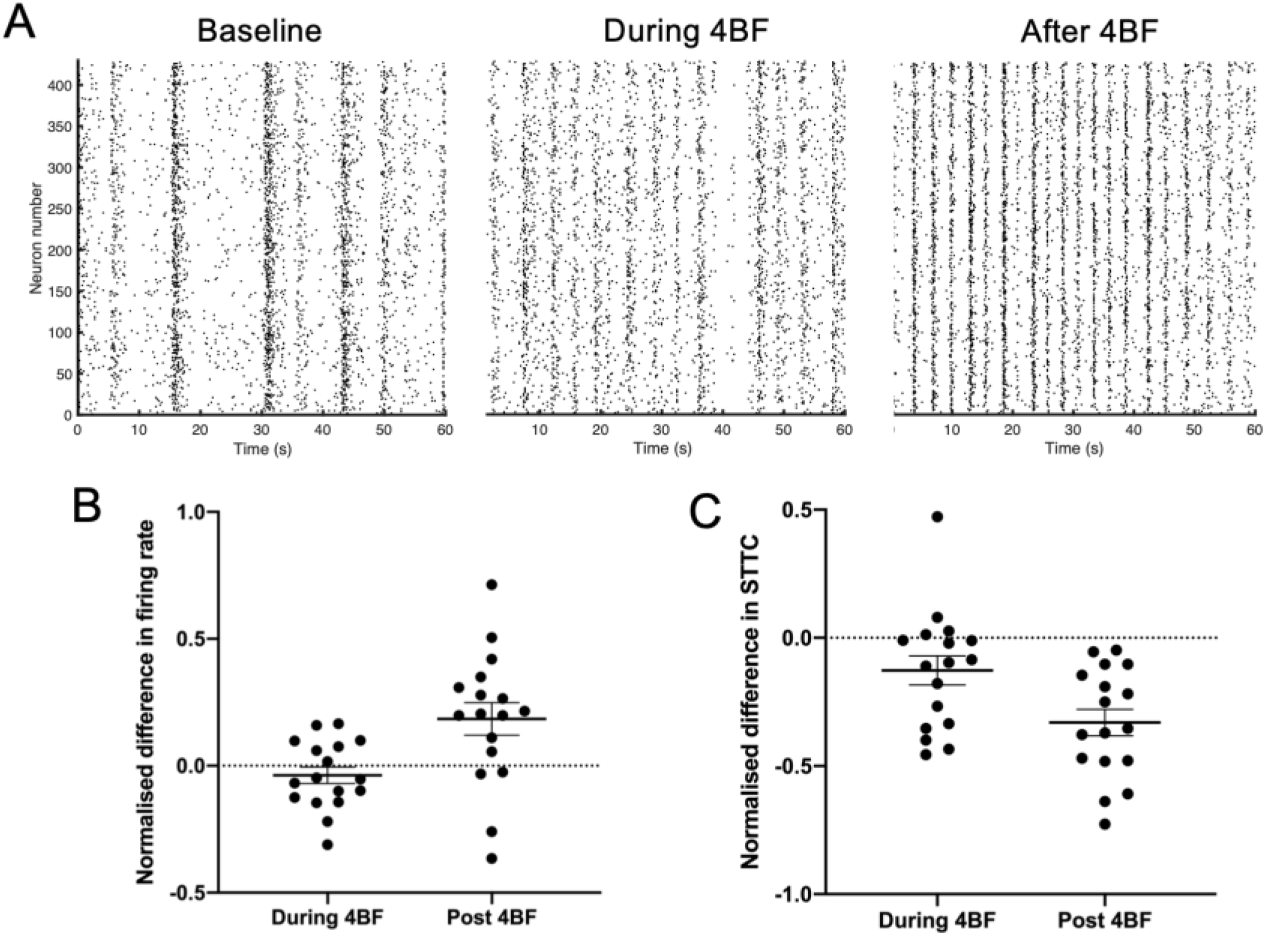
Neurons exhibit increased firing rate, but decreased connectivity after 4BF treatment. **A)** Representative raster plots of a culture at baseline and 3 hours after being treated with 4BF. **B)** Quantification of firing rates during and after 4BF treatment, both are compared and normalised to firing rates for each culture at baseline. Normalised difference in firing rate during 4BF = −0.0373, SEM = 0.0326. Normalised difference in firing rate post 4BF = 0.185, SEM = 0.0639. n = 17 for both. **C)** Quantification of STTC during and after 4BF treatment, both are compared and normalised to the STTC for each culture at baseline. Normalised difference in STTC during 4BF = −0.127, SEM = 0.0569. Normalised difference in STTC post 4BF = −0.330, SEM = 0.0518. n = 17 for both.

**Figure 3.**
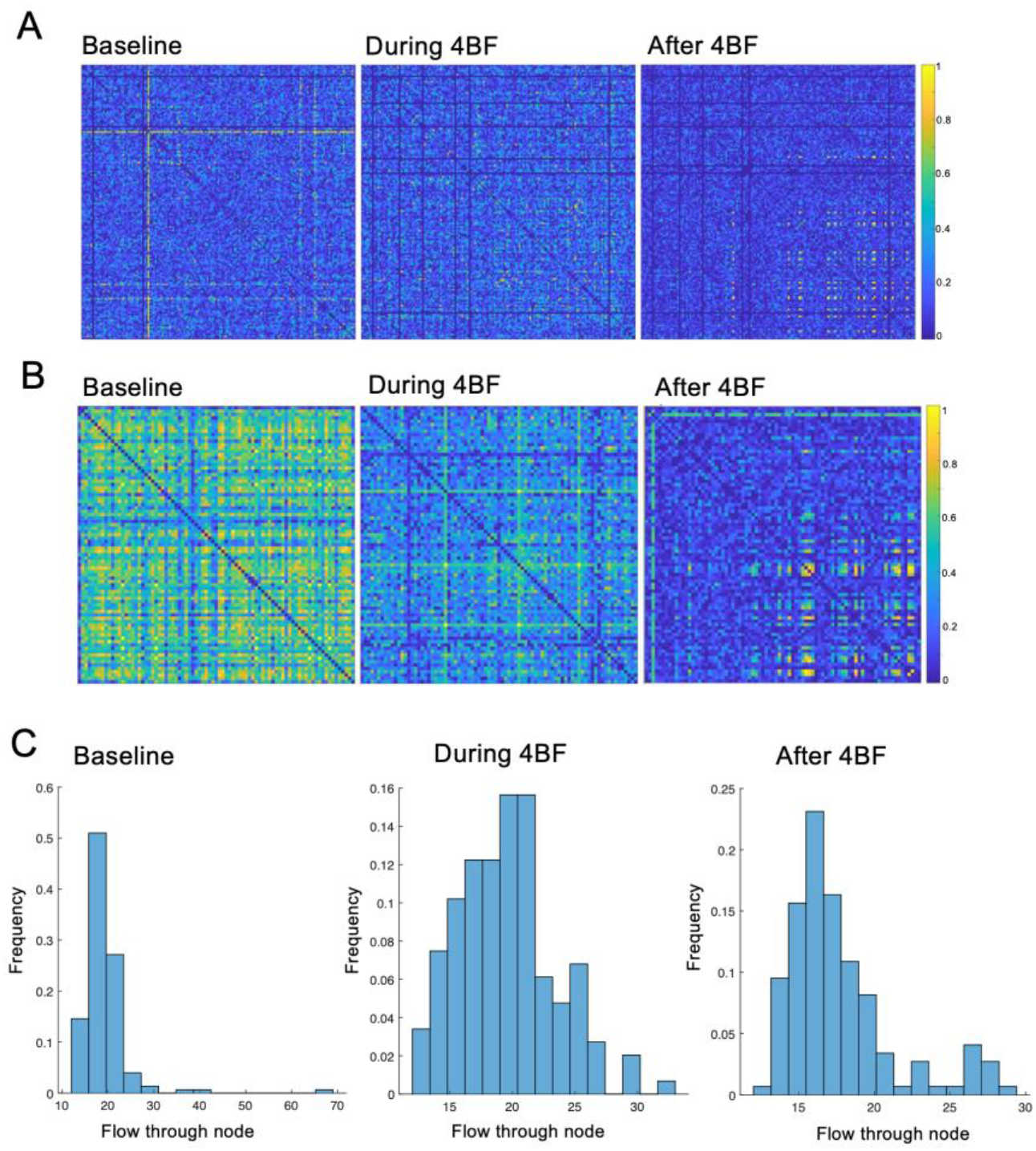
Refinement of connectivity after 4BF. **A)** A culture showing an initial state of low connectivity, with refinement of connectivity after 4BF. **B)** A culture showing an initial state of high connectivity with refinement of connectivity after 4BF. **C)** Histogram of total flow (sum of all connection strengths through each node) at baseline, and during and post 4BF for a representative culture. There is a right shift in the distribution with a higher proportion of connectivity in the culture being of higher value post 4BF.

### Refinement of network connectivity by 4BF

To further investigate the distribution of functional connectivity in the neuronal cultures, we examined the connectivity of each neuronal pair from the connectivity matrix and found that although there was a general reduction in connectivity after 4BF (Fig. 2C), some cultures had subsets of neurons that were highly connected after 4BF. This happened independent of initial states of the network, in both highly connected cultures as well as networks with low initial connectivity (Fig. 3A, B). This refinement effect can also be seen in the histogram of total flow (sum of all connection strengths through each node) as seen in Fig. 3C, where there is a small peak at the highly connected strengths (as shown in red box).

To better quantify this refinement in network connectivity, it is necessary to specifically identify the strongly connected neurons, characterized by high connection weights. However, inter-culture comparisons are not ideal, as the threshold for highly connected nodes can vary among cultures, and threshold changes affect the determination of hub neurons. Using a standardised threshold for all cultures to determine the refinement effect is therefore not ideal. To remedy this problem, we employed a previously developed measure named ICD (Scheinost 2012), which eliminates the need for setting arbitrary thresholds, and allows all of the connectivity information to be captured with only a few parameters. Fig. 3C shows the degree distribution of the connection strengths as histograms, before, during and after treatment with 4BF. Each degree distribution can give rise to a survival curve (Fig. 4A) by thresholding the degree distribution at different values (x-axis values). The values above the threshold degree were then integrated to obtain the normalised degree (y-axis values). This data was modelled as a stretched exponential, where the variance (α) and parameter (β) can be obtained from the estimated function (Fig. 4A). The ICD metric captures the distribution of connections and the ICD difference plot reveals changes in the connection strength distribution to effectively compare the connectivity of the network before and after 4BF treatment (Fig. 4B). The ICD difference curves for all the cultures examined can be seen in Fig. 4C. Quantification of these curves was subsequently performed based on the width, height and location of the peaks. For instance, a peak in ICD difference plot that is at a larger value of threshold indicates that there is a big change in connectivity strengths before and after 4BF, and the width of the peak in the ICD difference plots indicates how widespread the change in connectivity strengths is. As in the case of the refinements that were observed, there is not only a global reduction in connectivity strengths (corresponding to a larger threshold value at the peak) but also a wider distribution for the difference in connection strength values (larger width of the ICD difference plots). The width measure and the peak location were found to correlate well to whether the network has undergone refinement (Fig. 4D). However, given that the effect of refinement is verified visually using the connectivity matrices, these measure are unable to serve as objective standards to define if refinement has taken place.

**Figure 4.**
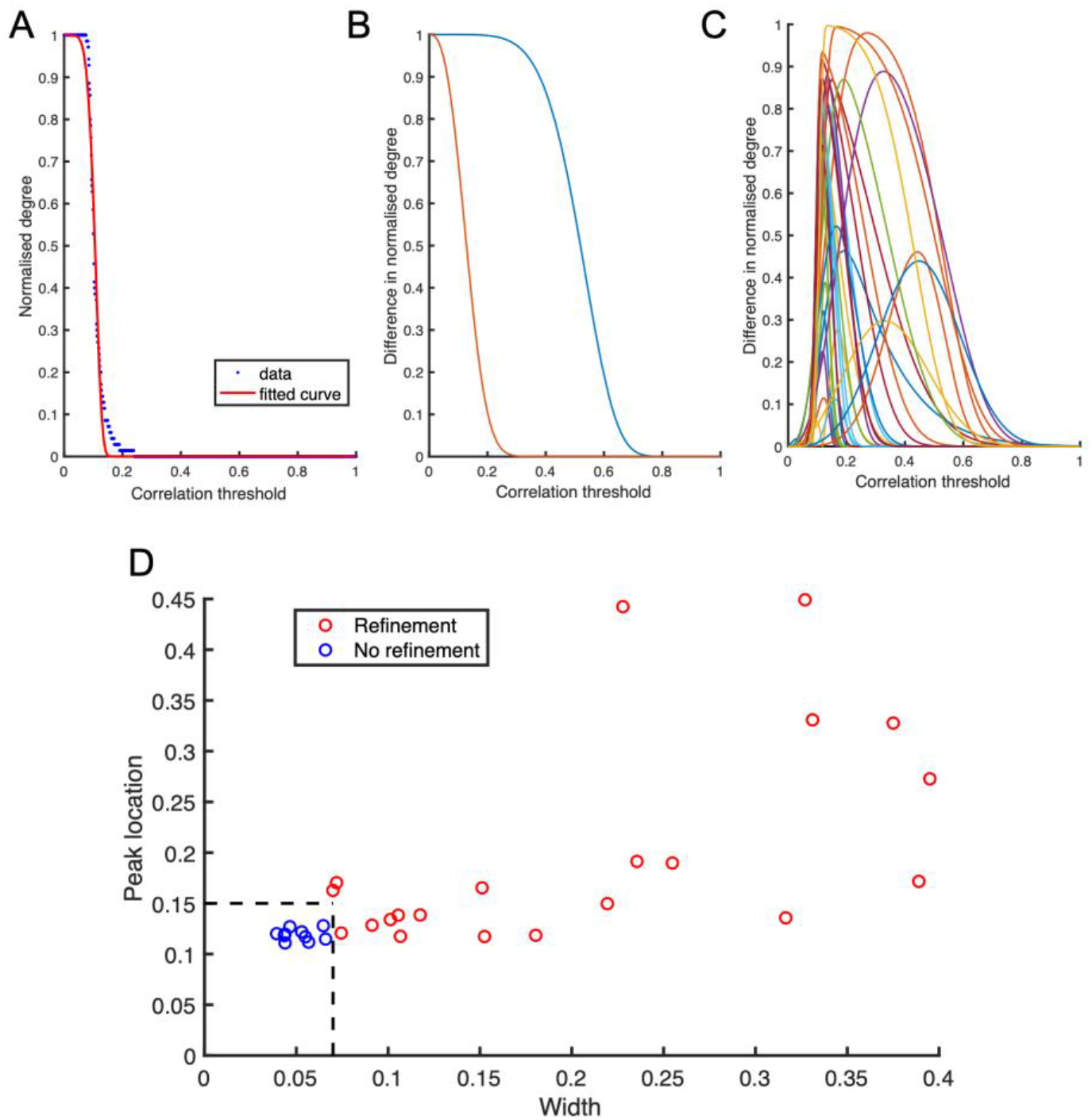
The ICD measure for describing neuronal connectivity. **A)** Survival curve obtained by thresholding the degree distribution at different values (x-axis values). The values above the threshold degree were integrated to obtain the normalised degree (y-axis values). Data at different values shown in blue, modelling of stretched exponential with variance (α) and parameter (β) (equation (2), methods), shown in red as the fitted curve. **B)** The ICD distribution at baseline (blue) and after 4BF treatment (orange) for a sample culture. **C)** ICD different plots for all cultures with each colour representing a different culture. The width, peak height and peak location of these curves can be obtained. **D)** Cultures that have not undergone refinement have low peak widths and peak locations (marked in blue ‘o’), n = 10. Cultures that have undergone refinement have large peak widths and peak locations (marked in red ‘o’), n = 21. Refinement was verified visually based on the connectivity matrices.

### Arc expression correlates with the final connectivity state and network refinement

To determine how overall network refinement may affects Arc expression, the percentage of Arc positive neurons was determined by immunofluorescent staining and categorically compared as in Fig 5. Cultures that express high levels of Arc seem to have undergone network refinement. To test this observation more robustly, a linear regression model was used to fit the data, with the final Arc expression percentage as the outcome. Of all input variables (including firing rate before and after 4BF, change in firing rate, connectivity indices before and after 4BF, change in connectivity index, and whether the network has undergone refinement), it was found that both the final connectivity index after 4BF treatment and whether the network has undergone refinement can best predict the level of Arc expression in the network (Fig. 5C, Table 1).

**Figure 5.**
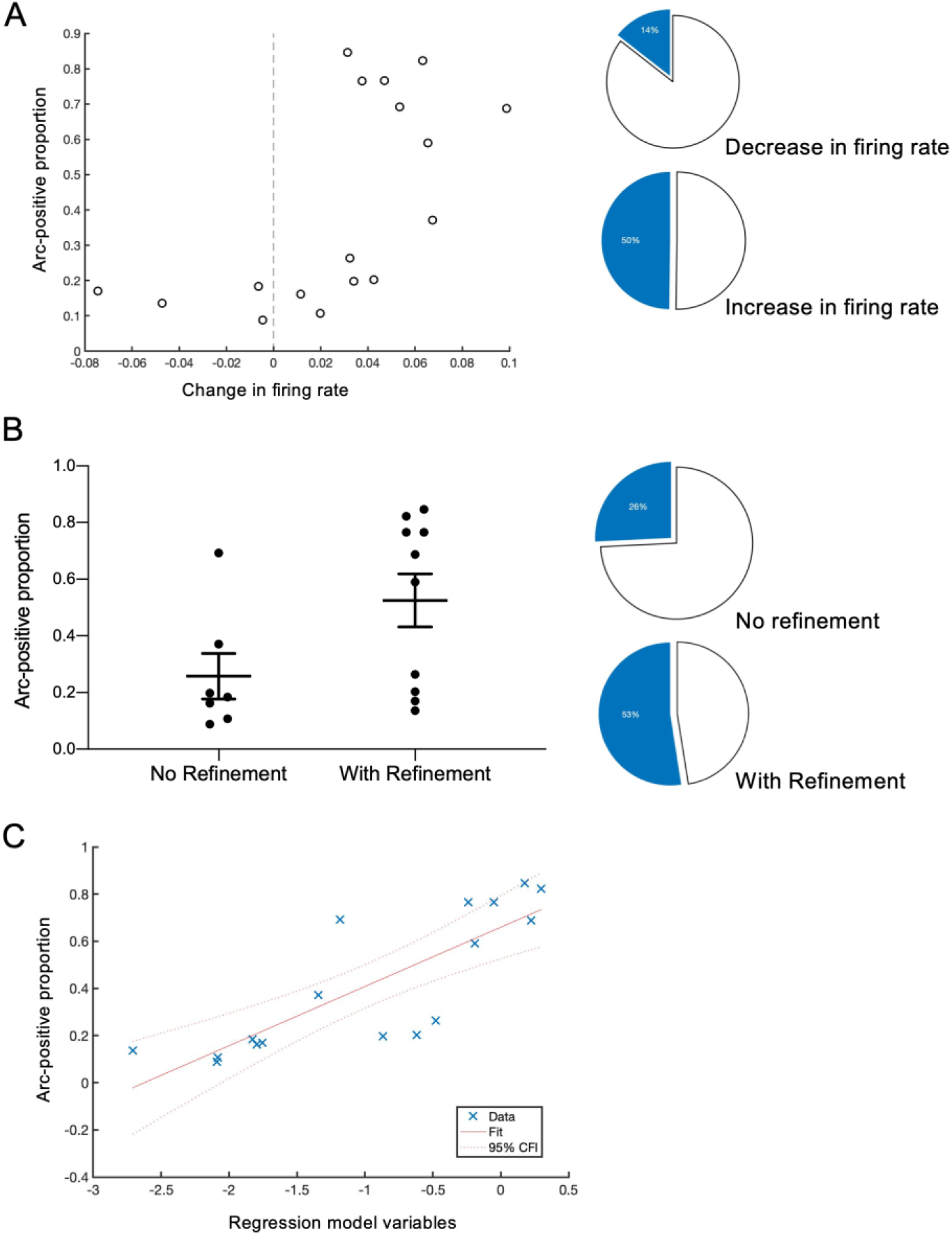
Arc expression and neuronal network properties. **A)** Graph showing change in firing rate after 4BF vs percentage of Arc expression in the culture. Side panel pie charts: a reduction in firing rate is associated with a low level of Arc expression (14%) while an increase in firing rate is associated with a higher level of Arc expression (50%). This was found from a total of n =17 cultures. **B)** Cultures with no refinement have a lower level of Arc expression as compared to cultures with refinement. Arc positive proportion in cultures with no refinement = 0.258, SEM = 0.0803, n = 7. Arc positive proportion in cultures with refinement = 0.525, SEM = 0.0935, n = 10. This can also be represented with pie charts: cultures with no refinement have an average of 26% Arc expression while those that have undergone refinement have a higher level of Arc expression (53%). **C)** Linear regression model with stepwise addition of factors connectivity (STTC values), firing rate, state of refinement (binary factor). It was found that the final STTC value together with the state of refinement are most able to explain the level of Arc expression after 4BF.

**Table 1.** Linear regression model variables and statistics

|  | Estimate | SE | tStat | pValue |
| --- | --- | --- | --- | --- |
| Final connectivity index | -0.02691 | 0.0060793 | -4.4265 | 0.0005748 |
| Refinement status | 0.24986 | 0.08757 | 2.8533 | 0.012766 |

### Arc positive cells are selectively strengthened and refined

To determine if there is any relationship between expression of Arc and functional connectivity, we specifically examined the STTC indices of Arc positive neurons in each culture and those that do not express Arc (only labelled with the neuronal marker NeuN). Results indicate that Arc positive neurons in each culture exhibit higher connectivity than other non-Arc neurons (Fig. 6A, B). This is the case at baseline, during and also after 4BF treatment (Fig. 6A). Cultures tend to have an overall reduction in connectivity after 4BF treatment (Fig. 6C). However, by separating the analysis into Arc and non-Arc neurons, it can be seen that for some cultures, the Arc-positive neuronal population actually increased their connectivity strength after 4BF (Fig. 6B). Specifically, some cultures which exhibited lower initial connectivity increased in connectivity only in the Arc-positive population but not in the Arc negative population (Fig. 6B, C). This selective increase can also be seen from the connectivity matrix, a representative one being shown in Fig. 6D.

**Figure 6.**
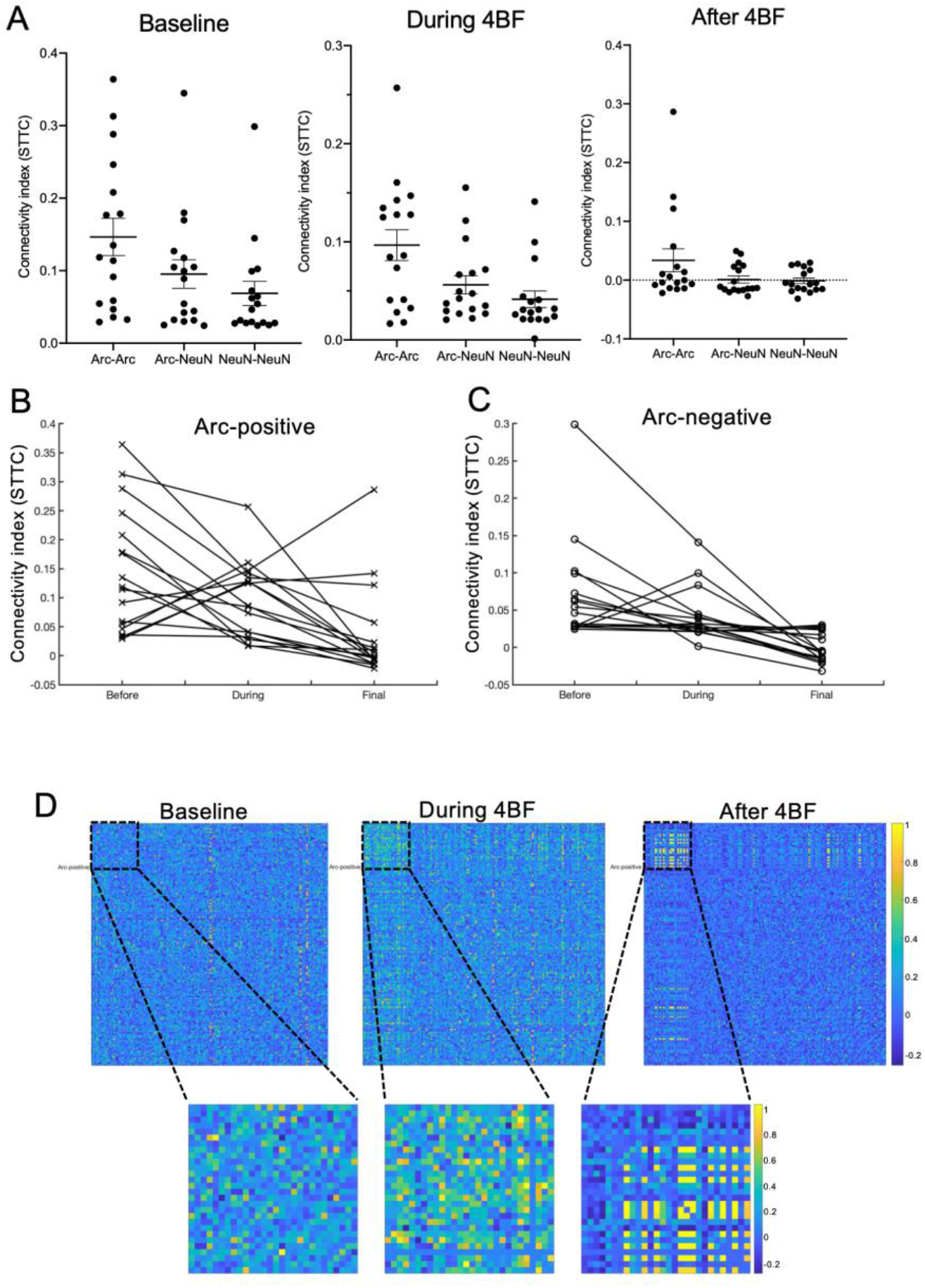
Difference between Arc-positive and Arc-negative neurons. **A)** Connectivity of Arc positive neurons in each culture and those that do not have Arc. Arc positive neurons in each culture exhibit higher connectivity than other non-Arc neurons at baseline, during and also after 4BF treatment. Mean connectivity index of Arc-Arc connections at baseline = 0.147, SEM = 0.0258. Mean connectivity index of Arc-NeuN connections at baseline = 0.0954, SEM = 0.0197. Mean connectivity index of NeuN-NeuN connections at baseline = 0.0686, SEM = 0.0166. Mean connectivity index of Arc-Arc connections during 4BF = 0.0967, SEM = 0.0157. Mean connectivity index of Arc-NeuN connections during 4BF = 0.0562, SEM = 0.00928. Mean connectivity index of NeuN-NeuN connections during 4BF = 0.0415, SEM = 0.00844. Mean connectivity index of Arc-Arc connections after 4BF = 0.0336, SEM = 0.0195. Mean connectivity index of Arc-NeuN connections after 4BF = 0.00108, SEM = 0.00598. Mean connectivity index of NeuN-NeuN connections after 4BF = −0.00127, SEM = 0.00474. A total of n = 17 cultures were examined. **B)** Some of the Arc-positive neuronal population increased in connectivity after 4BF. These were cultures which exhibited lower initial connectivity. **C)** Such increase is not seen in the Arc-negative population. D) Representative connectivity matrices of a culture at baseline, during and after 4BF treatment, showing selective increase and refinement of connectivity in the Arc-positive cells.

To investigate the effects of 4BF on connectivity especially in the Arc positive neuronal population, each individual connection between neurons was examined. These connections can be grouped into connections within the Arc-positive populations (that is the connections from Arc-positive neurons to Arc-positive neurons), connections between Arc-positive neurons and Arc-negative neurons, and connections within the Arc-negative neurons. The connection weights of these groups can be represented as a histogram as shown in Fig. 7A. It can be seen that the connections amongst Arc-positive neurons are stronger than the connections between Arc and non-Arc and within non-Arc neurons at baseline. This means that neurons that are initially well-connected to each other are more likely to express Arc after the 4BF treatment. Subsequently, after 4BF treatment, the overall connection strengths for all groups are reduced, but there is a small number of highly-connected strengths remaining, especially from the Arc-Arc group (Fig. 7A, right panel insert). To examine the change for each connection after 4BF treatment, the connection strength for each connection at baseline was plotted against its connection strength after 4BF (Fig. 7B). This analysis revealed that a large number of connections are depressed (downregulated in strength) after 4BF treatment (Fig. 7B, depressed connections box, right panel insert). However, a subpopulation of connections that are potentiated (upregulated in strength) after 4BF treatment (Fig. 7B, potentiated connections box, left panel insert). To determine the proportion of Arc-Arc connections among the potentiated versus depressed connections, the percentages of Arc-Arc connections were calculated as a function of the final (for potentiated) or initial (for depressed) connectivity index (labelled here as the connection threshold), as seen in Fig. 7C. This reveals that there is a large proportion of Arc-Arc connections which have very high connectivity after the 4BF treatment, which were initially weak connections. This therefore supports the hypothesis that a subset of connections between Arc-positive neurons are selectively strengthened after 4BF treatment.

**Figure 7.**
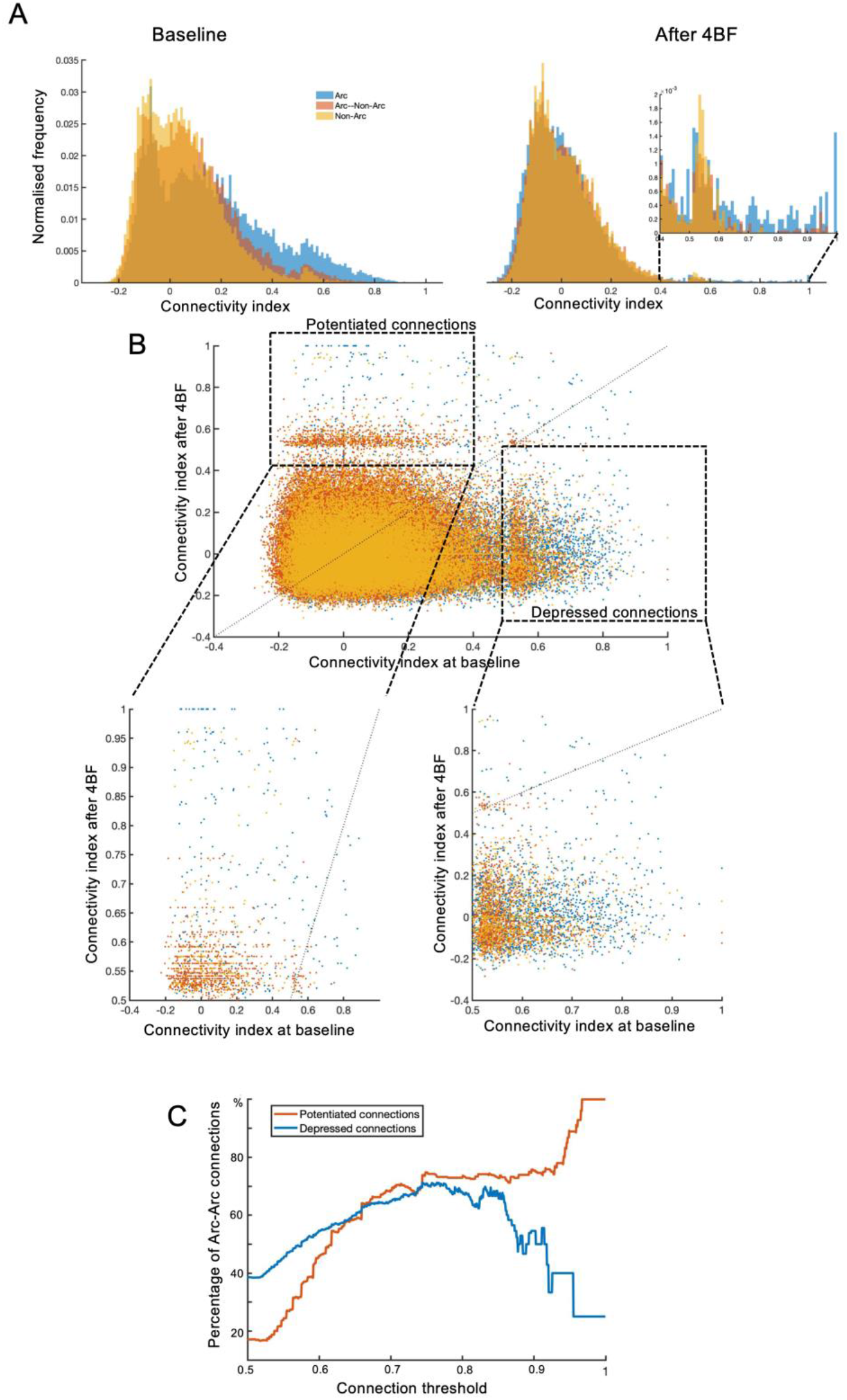
Difference between connections containing Arc-positive neurons and those that do not. **A)** Connections between pairs of neurons are grouped: Arc to Arc (blue), Arc to non-Arc (orange), and non-Arc to non-Arc (yellow). The distribution of connection strengths according to their groups (labelled in different colours) are shown in the histograms, with the right panel being the connection strengths between all pairs of neurons at baseline and the left panel being the connection strengths between all pairs of neurons after 4BF treatment. The x-axis represents the range of connection strengths, which are mostly between the values of −0.2 to 1. The y-axis represents the frequency at which each connection strength interval occurs, with a peak at around zero. This indicates that most of the connection strengths are fairly weak. The insert on the left panel is a zoomed-in view of the higher strength connections between 0.4 to 1. This shows that the highest connection strengths among pairs of neurons after 4BF treatment occur between Arc-positive and Arc-positive neurons. **B)** Scatter plot of connection weights before 4BF (at baseline) and after 4BF treatment. Left dotted box indicates connections that are weak initially that are strengthened after 4BF treatment (potentiated connections), whereas right dotted box marks out connections that are initially strong that become weakened after 4BF treatment (depressed connections). The lower left and right panels are zoomed in views of potentiated and depressed connections respectively. **C)** The percentage of Arc-positive to Arc-positive connections is determined by counting the numbers of Arc-Arc connections over the total number of connections at each connectivity threshold. Potentiated connections are those that have initial connectivity indices between the values of 0 and 0.5 and final values between 0.5 and 1 while depressed connections are defined as those that are initially weak between 0 and 0.5 and final values between 0.5 and 1. The x-axis represents the range of connection thresholds above which the percentage Arc-Arc connections are evaluated; for potentiated connections, the threshold is based on the final connectivity index while for depressed connections the thresholds are determined based on the initial connectivity index.

### Arc positive neurons are located physically closer to each other

To examine if there is any relationship between the type of connection (within Arc, between Arc and non-Arc and within non-Arc neurons) and the physical distances of these connections between the neurons, the coordinates of the neurons were obtained. Distances between each pair of neurons were calculated as described in the Materials and Methods section. Fig. 8A shows the values of distances between pairs of neurons for each type of connection. We found that the connections within Arc-positive neurons are generally shorter than the connections between Arc and non-Arc and also within the non-Arc neuron pairs. This means that Arc-positive neurons are likely to be closely located in the vicinity of each other and fewer long-range connections are found within the Arc-positive group. To determine if there is any correlation between the connection strength and physical distance between each pair of neurons, we examined the correlation between distances between pairs of neurons and their connectivity both at baseline (Fig. 8B, left panel) and after 4BF (Fig. 8B, right panel). It can be seen that both at baseline and after 4BF, the distribution of distance versus connectivity does not change much; most of the distances are short-ranged, with a few long-ranged connections that extend up to 2.4 mm. These long-range connections tend to be rather weak with connectivity indices around zero. This is the case both at baseline and also after 4BF treatment. The zoomed-in insert on the right panel of Fig. 8B shows that strong connections after 4BF treatment are likely to be short-range connections between pairs of neurons of which at least one is Arc-positive. This means that the strong connections between Arc-positive neurons and Arc-negative neurons tend to be short-range connections (Fig. 8C, strong connections between Arc and non-Arc neurons marked out with arrows). To determine if there is any difference in physical distance between potentiated versus depressed connections, a similar analysis was done as in the previous section. There is a general down sloping trend for both types of connections, showing that strongly modulated connections tend to be short-ranged (Fig. 8D).

**Figure 8.**
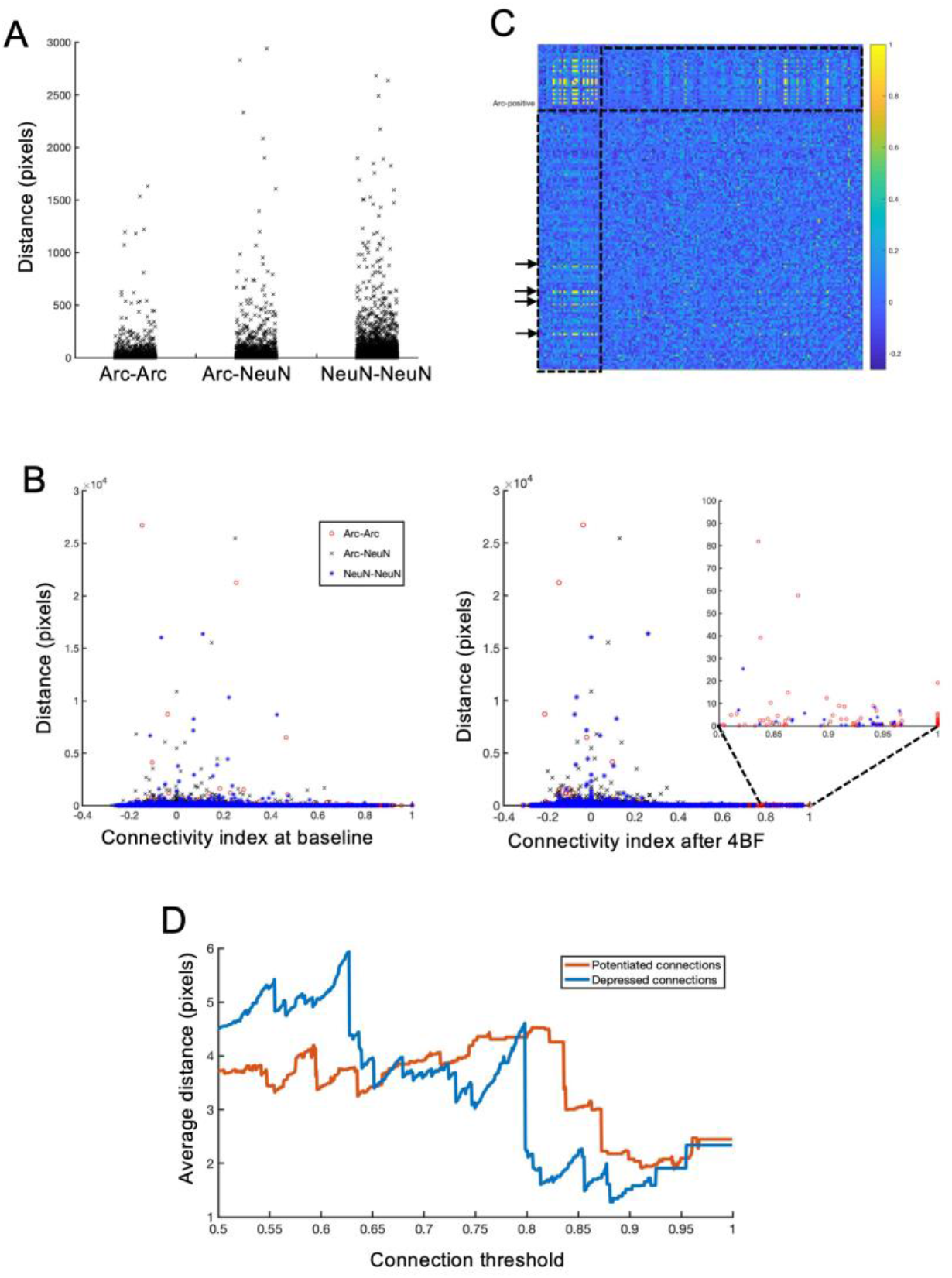
Distance measures of different types of connections. **A)** Distances between pairs of neurons for each type of connection. Arc-Arc: Arc-positive neurons with Arc positive neurons. Arc-NeuN: Arc-positive neurons with Arc-negative (non-Arc) neurons. NeuN-NeuN: Arc-negative neurons with Arc-negative neurons. **B)** Distances between pairs of neurons vs connectivity index at baseline (left panel) and after 4BF (right panel). Different types of connections marked with separate types of markers (see legend). Zoomed-in insert on the right panel shows the same graph with connection values between 0.8 and 1. **C)** Representative connectivity matrix after 4BF treatment (as in Figure 4.3D). Dotted boxes show the connectivity profile between Arc-positive and Arc-negative neurons, with arrows indicating strong connections with high indices. **D)** Potentiated connections are those that have initial connectivity indices between the values of 0 and 0.5 and final values between 0.5 and 1 while depressed connections are defined as those that are initially weak between 0 and 0.5 and final values between 0.5 and 1. The x-axis represents the range of connection thresholds above which the percentage Arc-Arc connections are evaluated; for potentiated connections, the threshold is based on the final connectivity index while for depressed connections the thresholds are determined based on the initial connectivity index. The y-axis shows values of average distances of all connections above the threshold value in x. Distance measured in pixels, with each pixel corresponding to 0.8 μm.

### Arc positive cells are mostly a subset of c-Fos cells

Arc and c-Fos are both well-studied IEGs and have been widely used in many settings as markers of memory engram neurons. However, not many studies have been done to determine the relationship of their expression and if they are expressed in different or similar populations of neurons after neuronal activation. To investigate the effects of 4BF treatment on the expression of both Arc and c-Fos *in vitro,* the same drug treatment protocol was used and the expression levels of both these proteins were assessed simultaneously using immunofluorescence staining. By co-labelling both Arc and c-Fos, it was found that most of the Arc expressing cells were also c-Fos-positive. A threshold is selected as described in the methods, and the numbers of Arc and/or c-Fos positive neurons can be quantified and their proportions determined across several cultures. The majority of neurons are positive for both Arc and c-Fos, with some that are only positive for c-Fos and only very few cells that are only positive for Arc (Fig. 9A). The pie chart in Fig. 9B shows that an average of 16.2% of all cells in the culture express Arc and/or c-Fos after 4BF treatment. Among the Arc-positive neurons, 82.7% of these are also c-Fos-positive (Fig. 9B).

**Figure 9.**
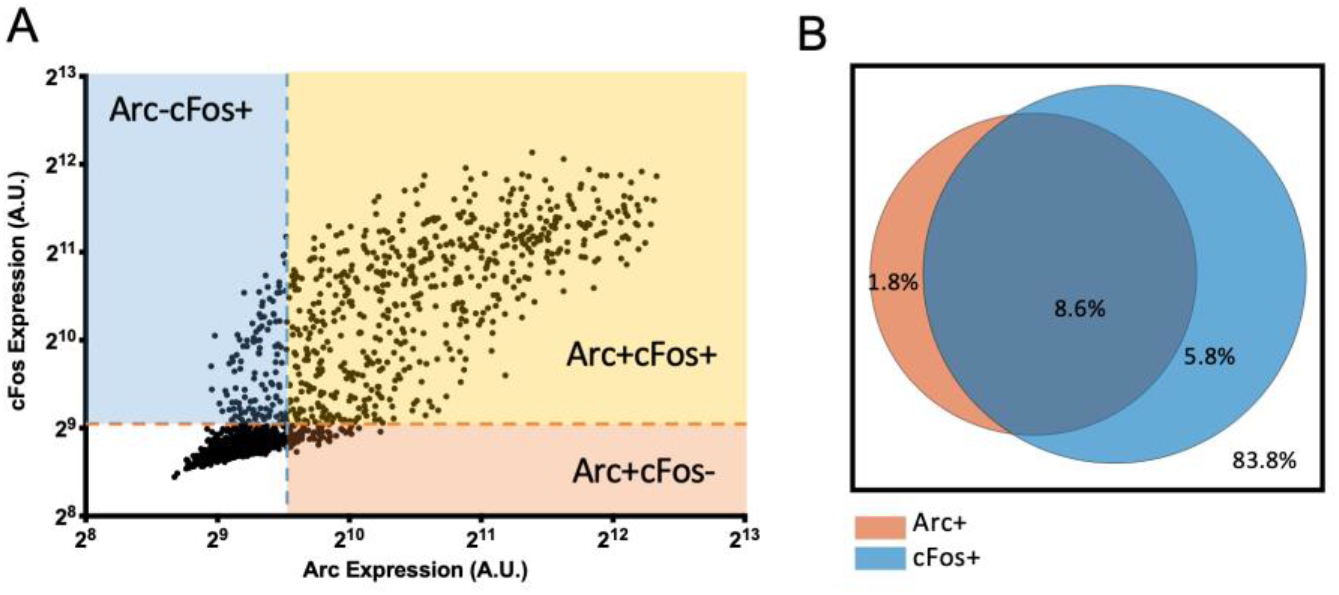
Arc and c-Fos expression after 4BF. **A)** Expression of Arc and c-Fos in individual neurons from a single frame. This shows the majority of the cells are both Arc and c-Fos positive, with a good number of c-Fos only expressing cells but few Arc only expressing neurons. **B)** Pie chart with Arc and/or c-Fos-positive percentages after 4BF treatment.

### Arc and c-Fos expression as a combinatorial code for connection strength pruning

Based on the expression profile of Arc and c-Fos, the cells can be grouped into Arc**+**cFos**+**, Arc**+**cFos**-**, Arc**-**cFos**+**, and Arc**-**cFos**-**. The intra group and across group connections between each pair of neurons can thus be classified into ten different sets. To examine the change across time after 4BF treatment on each set of connections, probability density distributions of each set of connections is plotted using a non-kernel density estimation method (Fig. 10A). The intra-group connection sets (Fig.10A, subplots I, V, VIII, X) are also plotted on the same graphs in Fig. 10B to highlight the differences across different intra-group sets. Results show that double positive neuronal connections are the strongest prior to 4BF treatment (Fig. 10B, left panel). The connections between neurons expressing only Arc but not c-Fos to other neurons were specifically down-regulated (Fig. 10A, subplots II, V, VII, Fig. 10B). Interestingly, the c-Fos-only connections were not changed much after 4BF (Fig.10A, subplots III, VI, VIII, IX). From these results, there seem to be a combinatorial code for regulating the strength of connections after 4BF as coded by the expression of Arc and/or c-Fos.

**Figure 10.**
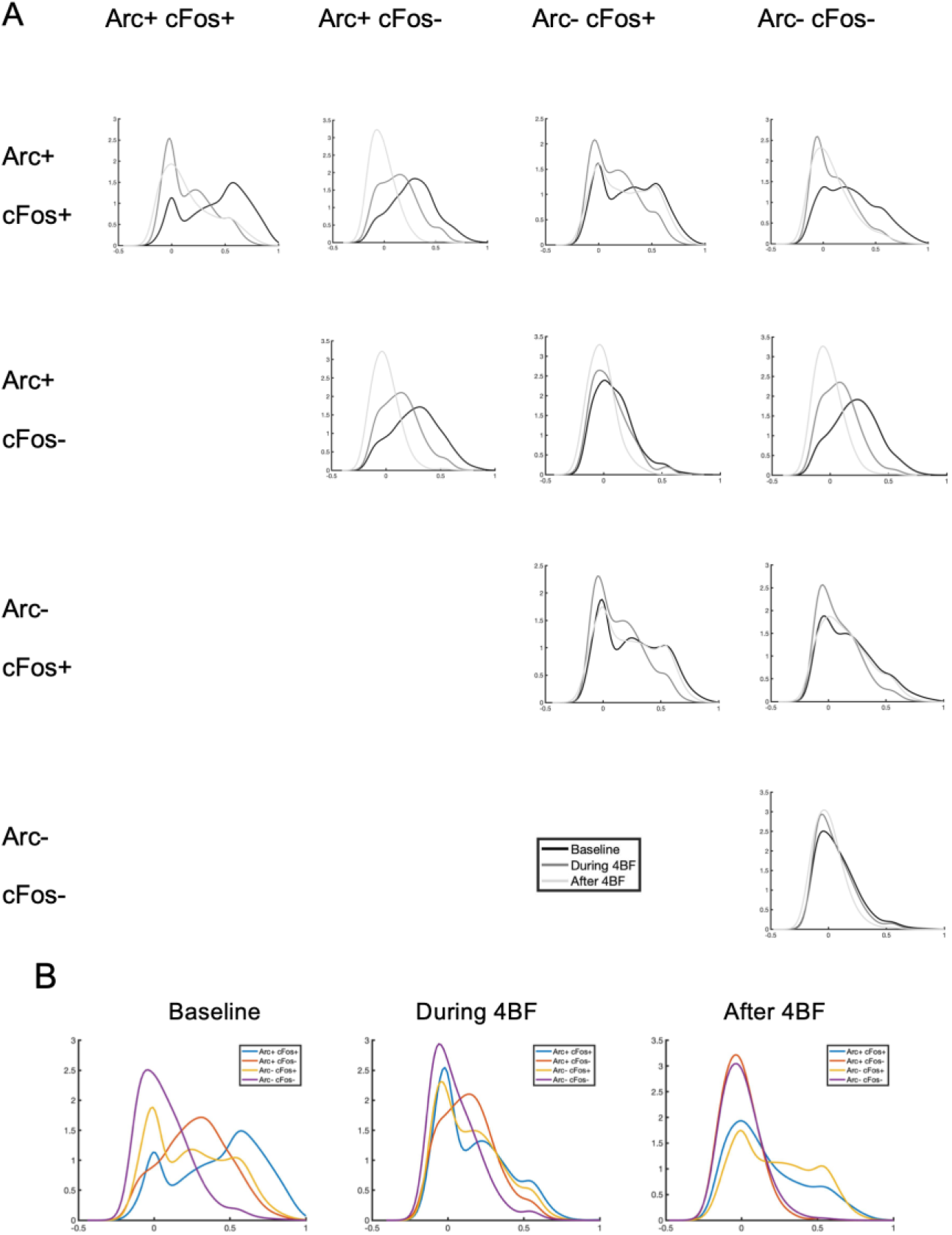
Probability density plots of connection strength distributions of different populations of neurons. (x-axis represents the connection strengths and y-axis represents frequency of the strengths for all plots.) **A)** Probability density plots of connection strength distributions between different groups of neurons that express Arc and/or c-Fos or none of these markers. Given that there are four groups of cells depending on the expression status (as listed in the top row and similarly in the first column), there are ten different sets of connections between these four groups. The strengths of connections between these 10 sets are non-overlapping and are represented in the subplots I to X. The row and column headers are labelled with the identity of the groups from which the connections are between. **B)** The intra-group connection sets (subplots along the diagonal, I, V, VIII and X) are extracted and plotted using different colours on the same graph for baseline, during 4BF treatment and after 4BF treatment.

## DISCUSSION

We have shown that the chemical induction of LTP using 4BF in cultured hippocampal neuronal networks led to a global reduction in network connectivity while selected connections are strengthened. This refinement effect correlates specifically with Arc expression in the network, with Arc-positive neurons having connections that are selectively strengthened. The expression patterns of IEGs Arc and c-Fos were shown to be mostly overlapping, but Arc is more selectively expressed than c-Fos post 4BF. Arc and c-Fos were also shown to act together in encoding information about connection strength pruning.

### Network refinement may be important for engram specification

The potentiation effects of LTP have been well-studied on the cellular level, with the measurable end result usually being increases in excitatory postsynaptic potentials (EPSPs) or changes in spine size. However, few studies have investigated the effects of changes in neuronal firing behaviour (Marder 2004). One such study by Chiappalone et al. (Chiappalone 2008) using neuronal cultures grown on MEAs found that a tetanus stimulus can induce an overall decrease in effective connectivity, similar to what we have reported here, although the same stimulus has been previously shown to induce potentiation of specific pathways (Jimbo 1999). It has long been thought that plasticity mechanisms cause networks to destabilise (Abbott 2000), and many cellular and network mechanisms, generally termed as homeostatic plasticity, have been discovered to push their state back to baseline (Turrigiano 2012). For instance, as shown in the seminal experiments of Turrigiano’s group, when the GABA receptor antagonist bicuculline was added acutely to cultured neurons, it produced an increase in firing rate in the short term but a return to baseline values in the long term (Turrigiano 1998). This effect is also known as synaptic scaling. Among the few cultures that had a decrease in firing rate in response to 4BF, all of them had high initial connectivity states, in accordance with homeostatic plasticity. However, there is a dissociation between the changes in firing rate and connectivity following 4BF treatment. Although most of the cultures showed an increase in firing rate, there is also a decrease in connectivity in all the cultures. This may signal that firing rate variations and connectivity changes take place on a different time scale. Homeostatic synaptic scaling that is required for firing rate modifications in response to prolonged neuronal activity is thought to involve the differential expression of AMPA receptor subunits (Goold 2010) and a reduction of AMPA receptors at the synapse (Seeburg 2008). Interestingly, Arc is known to mediate synaptic scaling of AMPA receptors in chronic inactivity but not in chronic activity (Shepherd 2006). The synaptic scaling that results from chronic activity is commonly observed at 48 hours after stimulus onset (Turrigiano 1998). The changes reported here on network connectivity take place in a shorter time frame, around 6 to 8 hours after 4BF is applied, and therefore may be different from the usual mechanisms known in synaptic scaling. This is supported by the fact that firing rates in most cultures still showed an increase at this time frame. Therefore, this global downregulation of network connectivity after 4BF treatment may occur via mechanism that is distinct from synaptic scaling.

It is known that synaptic plasticity can reach saturation, above which no further learning can occur (Moser 1998). Thus, down regulation of connectivity in response to prolonged activity is necessary for the network to continue coding useful information. Besides cell autonomous mechanism of achieving homeostasis, networks are known maintain stability through coordinated inter-cellular interactions (Maffei 2009). One feature worthy of mentioning is the criticality of neuronal networks. Criticality is a network state that maximises information processing capacities by having high level of responsiveness to external stimuli (Rocha 2018). Using neural networks *in silico,* it has been demonstrated previously that plasticity mechanisms can tune a network to criticality (Rubinov 2011). Effects of 4BF could push the network to criticality and possibly to a supercritical state, where the threshold of activation is high and thus reflects a global down-regulation of connectivity. This is in line with the effects of refinement seem in the connectivity patterns of the networks after 4BF. Adjustments in the network threshold are dynamic and could take place at the same time when plasticity mechanisms are in play. Such an ongoing state change for plasticity has been previously proposed and is part of the consideration of network homeostasis from an energy usage point of view, known as the “energy homeostasis principle” (Vergara 2019). These lines of evidence and the results shown here support the idea that regulation of network connectivity is state-dependent and interacts with mechanisms of plasticity.

Pharmacological induction, such as the 4BF treatment used here, is often thought to be a strong inducing stimulus that may not be physiologically relevant. However, repeated activation of NMDA receptors is known to be important for memory consolidation (Wittenberg 2002). It has also been shown that during sleep, the majority of cortical synapses are up-regulated, supporting the hypothesis that slow oscillations observed during sleep potentiate synapses that were depressed due to persistent activities during the day (Timofeev 2001). The strong inducing stimulus of 4BF thus may mimic the day stimulus that results in depression of synaptic connections due to persistent activity, and the global downregulation therefore serves to highlight the connections that need to be consolidated further. Therefore, this refinement process also acts to increases the contrast of weights in the network and may thus contribute to the process of engram specification and be crucial for memory consolidation.

### Arc-positive neuronal connections exhibit increased functional plasticity

IEG expression is often used as a marker of neuronal activity. We showed that in networks where the effect of 4BF on network refinement is largest, there is correspondingly a high level of Arc expression. Subsequently, not all the neurons in the network express Arc, even though they might participate in the increased network firing. In particular, Arc expression is high in neurons that have connection strengths that are strongly modulated, and especially for potentiated connections. This is consistent with findings *in vivo* by Mahringer *et al* (Mahringer 2019) where they found that neurons which would undergo the strongest learning-related changes in activity would exhibit increased levels of IEG expression. 4BF used here may act as a strong plasticity inducing stimulus, which causes network bursting that potentiates selective connections while depressing most others. This requires plasticity changes in both directions, with up and down-regulation of connection strengths accordingly. Arc may be poised to be performing a role in this process as it has been shown to play a part in both LTP as well as long-term depression (LTD) (Bramham 2008). Arc appears to be present in neurons which have connections that are strongly modulated, either potentiated or depressed. The concurrent presence of c-Fos may aid to distinguish between the subsets that are to either undergo potentiation or depression. In all, these results suggest that Arc-positive neuronal connections exhibit increased functional plasticity and supports the idea that Arc expression is not primarily a marker of neural activity, but of experience-dependent plasticity.

### Expression of Arc may depend on network architecture

We have shown that the average physical distance between neurons that are either strongly potentiated or depressed after 4BF treatment tends to be relatively small and these connections tend to involve at least one Arc-positive neuron. Short-range connections at the brain network level are known to be strongly linked. For instance, in fMRI studies, it has been shown that the functional connectivity is strongest between regions of shortest physical distance and that connectivity strength decreases as the physical distance between brain regions increases (Salvador 2005, Bellec 2006). Similarly, from diffusion spectrum imaging it is known that the degree of white matter tract connectivity also decays with distance (Honey 2009). This pattern of functional connectivity variation with physical distance aims to minimise wiring cost and thus follows economical principle in circuit organisation. Such a pattern of connectivity may also be at play in the *in vitro* setting for individual connections between neurons. Given the nature of the 4BF stimulus, which acts to increase synchronised bursting of the network, all neurons are technically active, though only a selective subset go on to express IEGs such as Arc. It can be seen in this *in vitro* hippocampal network that neurons of these selected subsets tend to be located physically closer to each other, implying that strongly modulated Arc-positive neuronal connections are short-ranged. The expression of Arc, therefore, may depend on network dynamics, including the architecture of the network. Short-range connectivity at the brain level has been studied previously in many neuropsychiatric disorders, notably autism spectrum disorder and schizophrenia, where the former is associated with an increase and the latter with a reduction in short-range connections (Ouyang 2017). Interestingly, aberrant short-range connection changes have also been reported in Alzheimer’s disease (Carmeli 2014, Phillips 2016). Given that short-ranged connections that undergo plasticity changes involve expression of Arc, this may implicate Arc in activity-dependent disease pathogenesis.

### Arc and c-Fos combinatorial coding

It is known that activation of hippocampal neurons that express IEGs during learning episodes can drive learned behaviour while inactivation can lead to impaired recall and surpasses learned behaviour (Tonegawa 2015). The expression of IEGs has also been shown to not simply correlate with neuronal activity, but specifically with changes in activity that correlate with learning (Mahringer 2019). As seen from our results, Arc and c-Fos are expressed in subsets of neurons after 4BF induction, and connections that are strongly potentiated or depressed contain high percentages of Arc-positive neurons. Further analysis of the co-staining with c-Fos reveals that connections within the Arc**+**cFos**-** subgroup of neurons are mostly depressed. On the other hand, connections within the double positive (Arc**+**cFos**+**) neurons, the Arc**-**cFos**+** neurons, and connections between these two groups are strong. This supports the hypothesis that expression of Arc and c-Fos may act as a combinatorial code for distinguishing the subpopulations of neurons that undergo different types of activity-dependent plasticity. Arc and c-Fos act in combination to specify whether hippocampal neurons are involved in memory reactivation using a novel environment exposure protocol in mice (Jaeger 2018). This is in part due to the difference in expression dynamics of Arc and c-Fos after the stimulus, where both Arc and c-Fos are expressed more abundantly 1-hour post novel environment exposure, and highly overlap, but only Arc persists through 5 hours after the exposure. As such, doubly positive Arc**+**cFos**+** neurons are reactivated or newly activated upon re-exposure to the same context, whereas Arc**+**cFos**-** neurons are not reactivated upon exposure to a new context (Jaeger 2018). The prolonged presence of Arc post stimulation, compared to c-Fos more transient expression, is also found *in vitro* using pharmacological stimulation, which shows that some expression dynamics can be conserved even through different preparations. Thus, the changes in connection strengths between different groups of IEG-expressing neurons found here are potentially relevant to mechanisms of learning and memory. In the case of novel environment exposure, where the second exposure is to a new context, it would be necessary to down regulate the connections strengths previously strengthened in the first context, so that connections made in the second exposure can be more prominent. This down regulation would be for connections between Arc**+**cFos**-** neurons. On the other hand, the doubly labelled Arc**+**cFos**+** group has some connections down regulated, but most strengths are still high. This would be also in line with the novel environment exposure in which the new or re-exposure context would activate both c-Fos and Arc. Jaeger et al. did not specifically analyse the Arc**-**cFos**+** subset, for which we have shown that the connection strengths are high and remain mostly unchanged after the 4BF stimulus. This high c-Fos expression without Arc could code for on-going neuronal activation without the need for synaptic modulation - either increase or decrease in connection strength - and could point to a population of neurons that are activated during a behaviour or memory encoding process but are already well-connected and thus do not need further modulation. This also supports the idea that pre-existing network connectivity affects the formation of the engram and interacts with ongoing activity with differential expression of IEGs in response to neuronal activation.

### Dynamic interplay between network activity, IEG expression and plasticity

We made use of post calcium imaging immunostaining to characterize individual neurons and relate the expression of IEGs to the prior activity states of the neurons and subsequent changes in their connectivity. We believe that this is a major area that needs further research: the relationship between IEG expression, network activity and changes in plasticity. Conventionally, studies of neuronal activity and IEG expression are mostly based on the single cell level whereas network activity is recorded on the neuronal population level. There appears to be a lack in linking IEG expression and changes in plasticity on the cellular level with changes in network responses. Mahringer *et al* (Mahringer 2019) examined the expression of IEGs *in vivo* and linked it to the activity of neurons during specific tasks. This work examines responses from individual neurons and thus has the benefit of relating individual neuronal responses to specific stimulus presented to the animal. However, it did not address how the connections of the neuronal ensemble evolved during the process. In our study, we provided evidence from this missing link, although as an *in vitro* study, we are not able to link our finding directly to a certain behavioural output. Our results are in line with the hypothesis that IEG expression levels are correlated with changes in plasticity and go further to suggest that the pattern of expression also influences the network connectivity profile. To further probe the links between IEG expression and connectivity changes, specific stimulus patters selectively activating subsets of neurons could be used, for instance using optogenetic methods. This would provide a direct assessment of the causal relationship between changes in network activity and molecular changes within individual cells.

By making use of IEG-labelled engrams, it has been proposed that it is possible to introduce false memories in rodents (Ramirez 2014). Recent work has also independently shown that memory can be formed in the absence of experience (Vetere 2019). These studies were based on the prior knowledge of specific populations of cells or pathways that are activated during specific memory formation processes, and thus artificial stimulation of these pathways generated associativity between cellular networks leading to the formation of memory. However, the challenge remains to define specific populations of cells used in or pathways relevant for specific memory traces, especially without prior exposure and recording of the population of cells during the encoding process. Further complicating this is the ongoing interactions between IEG expression, mechanisms of plasticity and network activity. Therefore, an important step towards understanding learning and memory is to study how plasticity develops at the network level.

### Summary

We investigated the network changes that can occur after induction of LTP *in vitro* and found that chemical LTP induction by 4BF resulted in an emergent effect of refinement of the network connectivity. Arc expression correlates with effects of 4BF, particularly on network refinement, and Arc-positive neurons are selectively strengthened and refined. The architecture of the network may affect Arc expression as Arc positive neurons are located closer to each other in the network. We also examined the expression of IEGs Arc and c-Fos and found that Arc is more selectively expressed. These two IEGs also act together in coding information about connection strength pruning. With these results we have uncovered some interesting and important links between network activity, IEG expression and changes in connection strengths, which serve to further our understanding of learning and memory.

## REFERENCES

Abbott, L. F. and S. B. Nelson (2000). “Synaptic plasticity: taming the beast.” Nature neuroscience 3(11): 1178–1183.

Bellec, P., et al. (2006). “Identification of large-scale networks in the brain using fMRI.” Neuroimage 29(4): 1231–1243.

Bloomer, W. A., et al. (2008). “Arc/Arg3. 1 translation is controlled by convergent N-methyl-D-aspartate and Gs-coupled receptor signaling pathways.” Journal of Biological Chemistry 283(1): 582–592.

Bramham, C. R., et al. (2008). “The immediate early gene arc/arg3.1: regulation, mechanisms, and function.” J Neurosci 28(46): 11760–11767.

Carmeli, C., et al. (2014). “Structural covariance of superficial white matter in mild Alzheimer’s disease compared to normal aging.” Brain and behavior 4(5): 721–737.

Chiappalone, M., et al. (2008). “Network plasticity in cortical assemblies.” European Journal of Neuroscience 28(1): 221–237.

Cutts, C. S. and S. J. Eglen (2014). “Detecting pairwise correlations in spike trains: an objective comparison of methods and application to the study of retinal waves.” Journal of Neuroscience 34(43): 14288–14303.

Denny, C. A., et al. (2014). “Hippocampal memory traces are differentially modulated by experience, time, and adult neurogenesis.” Neuron 83(1): 189–201.

Gonzales, B. J., et al. (2019). “Subregion-specific rules govern the distribution of neuronal immediate-early gene induction.” Proceedings of the National Academy of Sciences: 201913658.

Goold, C. P. and R. A. Nicoll (2010). “Single-cell optogenetic excitation drives homeostatic synaptic depression.” Neuron 68(3): 512–528.

Hardingham, G. E., et al. (2001). “Nuclear calcium signaling controls CREB-mediated gene expression triggered by synaptic activity.” Nature neuroscience 4(3): 261–267.

Honey, C. J., et al. (2009). “Predicting human resting-state functional connectivity from structural connectivity.” Proceedings of the National Academy of Sciences 106(6): 2035–2040.

Jaeger, B. N., et al. (2018). “A novel environment-evoked transcriptional signature predicts reactivity in single dentate granule neurons.” Nat Commun 9(1): 3084.

Jimbo, Y., et al. (1999). “Simultaneous induction of pathway-specific potentiation and depression in networks of cortical neurons.” Biophysical journal 76(2): 670–678.

Kawashima, T., et al. (2014). “A new era for functional labeling of neurons: activity-dependent promoters have come of age.” Frontiers in neural circuits 8: 37.

Leung, H.-W., et al. (2019). “Arc Regulates Transcription of Genes for Plasticity, Excitability and Alzheimer’s Disease.” bioRxiv: 833988.

Liu, X., et al. (2012). “Optogenetic stimulation of a hippocampal engram activates fear memory recall.” Nature 484(7394): 381–385.

Maffei, A. and A. Fontanini (2009). “Network homeostasis: a matter of coordination.” Current opinion in neurobiology 19(2): 168–173.

Mahringer, D., et al. (2019). Expression of c-Fos and Arc in hippocampal region CA1 marks neurons that exhibit learning-related activity changes, bioRxiv.

Marder, C. P. and D. V. Buonomano (2004). “Timing and balance of inhibition enhance the effect of long-term potentiation on cell firing.” Journal of Neuroscience 24(40): 8873–8884.

Moser, E. I., et al. (1998). “Impaired spatial learning after saturation of long-term potentiation.” Science 281(5385): 2038–2042.

Nikolaienko, O., et al. (2018). “Arc protein: a flexible hub for synaptic plasticity and cognition.” Semin Cell Dev Biol 77: 33–42.

Oey, N. E., et al. (2015). “A Neuronal Activity-Dependent Dual Function Chromatin-Modifying Complex Regulates Arc Expression(1,2,3).” eNeuro 2(1).

Otmakhov, N., et al. (2004). “Forskolin-induced LTP in the CA1 hippocampal region is NMDA receptor dependent.” J Neurophysiol 91(5): 1955–1962.

Ouyang, M., et al. (2017). “Short-range connections in the developmental connectome during typical and atypical brain maturation.” Neuroscience & Biobehavioral Reviews 83: 109–122.

Patel, T. P., et al. (2015). “Automated quantification of neuronal networks and single-cell calcium dynamics using calcium imaging.” Journal of neuroscience methods 243: 26–38.

Phillips, O. R., et al. (2016). “The superficial white matter in Alzheimer’s disease.” Human brain mapping 37(4): 1321–1334.

Plath, N., et al. (2006). “Arc/Arg3.1 is essential for the consolidation of synaptic plasticity and memories.” Neuron 52(3): 437–444.

Poli, D., et al. (2016). “From functional to structural connectivity using partial correlation in neuronal assemblies.” Journal of neural engineering 13(2): 026023.

Ramirez, S., et al. (2014). “Identification and optogenetic manipulation of memory engrams in the hippocampus.” Frontiers in behavioral neuroscience 7: 226.

Rocha, R. P., et al. (2018). “Homeostatic plasticity and emergence of functional networks in a whole-brain model at criticality.” Scientific reports 8(1): 1–15.

Rubinov, M., et al. (2011). “Neurobiologically realistic determinants of self-organized criticality in networks of spiking neurons.” PLoS Comput Biol 7(6): e1002038.

Salvador, R., et al. (2005). “Undirected graphs of frequency-dependent functional connectivity in whole brain networks.” Philosophical Transactions of the Royal Society B: Biological Sciences 360(1457): 937–946.

Scheinost, D., et al. (2012). “The intrinsic connectivity distribution: a novel contrast measure reflecting voxel level functional connectivity.” Neuroimage 62(3): 1510–1519.

Seeburg, D. P. and M. Sheng (2008). “Activity-induced Polo-like kinase 2 is required for homeostatic plasticity of hippocampal neurons during epileptiform activity.” Journal of Neuroscience 28(26): 6583–6591.

Sheng, H. Z., et al. (1995). “Combinatorial expression of immediate early genes in single neurons.” Brain Res Mol Brain Res 30(2): 196–202.

Sheng, M. and M. E. Greenberg (1990). “The regulation and function of c-fos and other immediate early genes in the nervous system.” Neuron 4(4): 477–485.

Shepherd, J. D. and M. F. Bear (2011). “New views of Arc, a master regulator of synaptic plasticity.” Nat Neurosci 14(3): 279–284.

Shepherd, J. D., et al. (2006). “Arc/Arg3. 1 mediates homeostatic synaptic scaling of AMPA receptors.” Neuron 52(3): 475–484.

Timofeev, I., et al. (2001). “Disfacilitation and active inhibition in the neocortex during the natural sleep-wake cycle: an intracellular study.” Proceedings of the National Academy of Sciences 98(4): 1924–1929.

Tonegawa, S., et al. (2015). “Memory engram storage and retrieval.” Current opinion in neurobiology 35: 101–109.

Turrigiano, G. (2012). “Homeostatic synaptic plasticity: local and global mechanisms for stabilizing neuronal function.” Cold Spring Harbor perspectives in biology 4(1): a005736.

Turrigiano, G. G., et al. (1998). “Activity-dependent scaling of quantal amplitude in neocortical neurons.” Nature 391(6670): 892–896.

Vergara, R. C., et al. (2019). “The Energy Homeostasis Principle: Neuronal energy regulation drives local network dynamics generating behavior.” Frontiers in computational neuroscience 13: 49.

Vetere, G., et al. (2019). “Memory formation in the absence of experience.” Nature neuroscience 22(6): 933–940.

Vogelstein, J. T., et al. (2010). “Fast nonnegative deconvolution for spike train inference from population calcium imaging.” Journal of neurophysiology 104(6): 3691–3704.

Wee, C. L., et al. (2014). “Nuclear Arc Interacts with the Histone Acetyltransferase Tip60 to Modify H4K12 Acetylation(1,2,3).” eNeuro 1(1).

Wittenberg, G. M. and J. Z. Tsien (2002). “An emerging molecular and cellular framework for memory processing by the hippocampus.” Trends in neurosciences 25(10): 501–505.

Yap, E. L. and M. E. Greenberg (2018). “Activity-Regulated Transcription: Bridging the Gap between Neural Activity and Behavior.” Neuron 100(2): 330–348.

